# This is GlycoQL

**DOI:** 10.1101/2022.04.14.488348

**Authors:** Catherine Hayes, Vincenzo Daponte, Julien Mariethoz, Frederique Lisacek

**Affiliations:** Department of Computer Science, University of Geneva, Geneva, 1227, Switzerland; Proteome Informatics Group, SIB Swiss Institute of Bioinformatics, Geneva, 1211, Switzerland; Section of Biology, University of Geneva, Geneva, 1211, Switzerland

## Abstract

**Motivation:** We have previously designed and implemented a tree-based ontology to represent glycan structures with the aim of searching these structures with a glyco-driven syntax. This resulted in creating the GlySTreeM knowledge-base as a linchpin of the matching procedure and we now introduce a query language, called GlycoQL, for the actual implementation of a glycan structure search.

**Results:** The methodology is described and illustrated with a use-case focused on SARS-CoV-2 spike protein glycosylation. We show how to enhance site annotation with federated queries involving UniProt and GlyConnect, our glycoprotein database.

**Availability:** currently only available for reviewers at: https://beta.glyconnect.expasy.org/glycoql/

**Supplementary information:** Supplementary data are available at https://glyconnect.expasy.org/glystreem/wiki.

## 1 Introduction

In the last twenty years, bioinformatics resource interoperability has evolved from being a conceptual view and a difficult goal to achieve to becoming a concrete and frequent concern of database developers. A wide spread technological solution is provided by the semantic web and an increasing number of databases are now RDFized and accordingly, a broad range of triple stores is accessible on-line. With the deployment of the corresponding SPARQL endpoints, multiple data sources can be simultaneously searched and results aggregated to support data integration. Such an expansion of semantic web technologies is an opportunity for glycoinformatics to contribute glycobiology knowledge, otherwise considered too complex or too confusing, to the overall biology picture. The first step in understanding and modeling glycans lies in capturing their tree-like structure while coping with a high level of ambiguity generated by still non optimal experimental workflows. Several encoding schemes such as, IUPAC condensed (Sharon (1986)) KCF (Kotera *et al*. (2013)), GlycoCT (Herget *et al*. (2008)), WURCS (Tanaka *et al*. (2014)) have been proposed. While the linear summary of IUPAC sequences remains compelling for many bench glycoscientists, glycoinformaticians mainly rely on GlycoCT and WURCS. Actually, the latter are currently implemented in PubChem glycan entries (Kim *et al*. (2019)) and soon expected to be in ChEBI (Hastings *et al*. (2016)). Many thousands of glycan molecules are stored in GlyTouCan (Fujita *et al*. (2021)) the universal structure repository. Ideally, a fully characterised glycan is defined with a precise set of monosaccharides, each one linked to another with precise linkage details (anomericity, glycosidic bonds). In reality, many structures are provided with missing information. A mannose may not be distinguishable from a galactose as both are hexoses and only identified as such. Likewise, the precise carbon atoms involved in linkages are often unknown despite the importance of the distinction. In the end, GlyTouCan and other glycan databases collect redundant and ambiguous glycan entries with the potential to fully or partially match each other but no obvious means to ascertain it. Yet, glycan expression details are key to understanding cell-cell communication and other biological processes making structure comparison a necessary step. Figure 1 shows examples of the data status where five unevenly defined structures extracted from the GlyConnect database (Alocci *et al*. (2019)) are shown. The central structure is fully characterised and as such, found to be attached to 55 proteins in various tissues and species while the four other structures with the same monosaccharide composition and ambiguous to unknown linkages, are found specifically on one protein. The uniqueness of the four outliers is difficult to interpret. Note that all throughout this article, glycans are represented in the Symbol Nomenclature For Glycans (SNFG) widely accepted in the glycoscience community (Neelamegham *et al*. (2019)).

**Fig. 1:**
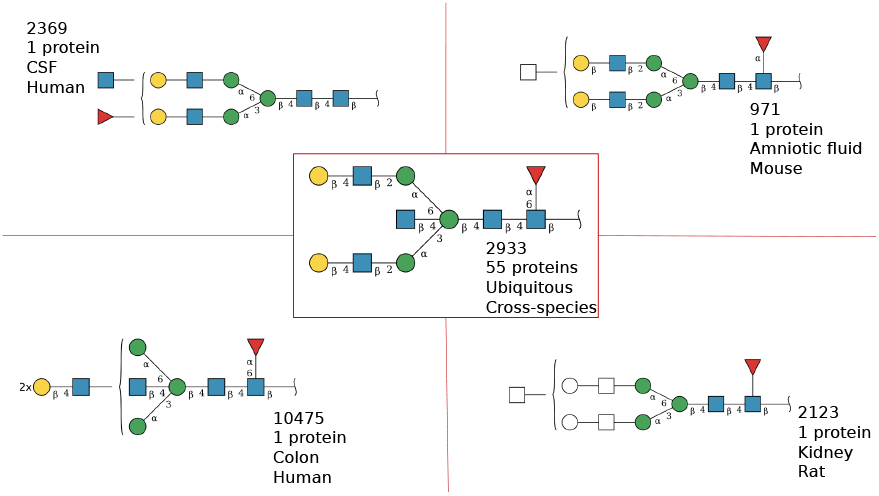
Spread and irreconcilable annotations due to the ambiguity of glycan structures. Each structure is depicted in SNFG and shown with its GlyConnect ID, number of proteins where it is attached, the tissue in which it is expressed and in which species.

In this context, (sub)structure search tools are needed for comparison purposes and further biological interpretation. A few have been proposed and implemented as a web tool relying on different encoding and models, e.g., GlyS3 (Alocci *et al*. (2015)), GlycoGlyph (https://glycotoolkit.com/glycoglyph, SugarDrawer (Tsuchiya *et al*. (2021). Originally defined with an RDF model, GlyS3 cannot properly handle structural ambiguity, while GlycoGlyph and SugarDrawer that are not RDF-based, allow for the drawing of ambiguous structures but with the purpose of retrieving an exact match from a database. In the end, neither of these tools can actually search a precise structure in a collection of ambiguous ones. To solve this problem and enable fuzzier matching from both the query and the result viewpoints, we recently designed a tree-based ontology to develop GlySTreeM, a glycan structure RDF knowledge base (Daponte *et al*. (2021)). GlySTreeM has proved an efficient tool to represent the structure of glycans and enable flexible searches within and across these structures. We now bring this initiative further by introducing a query language that takes advantage of the tree representation in order to search patterns. This requires a translation into a suitable query language such as SPARQL.

(Sub)structures to be queried may be complex, for example a type of core structure or an ambiguous motif. The effort needed to exploit the features of GlySTreeM and complete these queries can drive away scientists that are not familiar with semantic web technologies. To restrict this contingency and widen the availability of the GlySTreeM (sub)structure search, we developed a new SPARQL-inspired approach called GlycoQL that eases glycan (sub)structure queries based on a syntax recognised in glycoscience. The present article describes this new approach.

## 2 Approach

The aim of this work is to provide the means to consistently perform (sub)structure searches on the glycan structure triple store, GlySTreeM, Daponte *et al*. (2021) without expert knowledge of the SPARQL query language. The GlySTreeM pipeline is based on the GlycoCT encoding scheme, Herget *et al*. (2008), a widely adopted input and storage format for glycan structures. This choice was preserved for defining GlycoQL, but can be extended to other structure formats provided there is suitable alignment between the syntax and the underlying GlySTreeM ontology model.

The data-import algorithm used in GlySTreeM, takes GlycoCT strings and creates an RDF representation of the glycan tree structure. To search for a given (sub)structure, the query pattern should mimic this tree structure, in the form of a SPARQL query.

The GlySTreeM data-import algorithm was used as a starting point in the development of GlycoQL. The main difference between the output of the GlySTreeM pipeline, and that of GlycoQL is in the translation of undefined values in the GlycoCT strings. In GlySTreeM, undetermination is considered as a piece of knowledge. In the context of a search parameter there is a subtle but important difference; an undefined value in the search query would imply patterns that match any possible value. GlycoQL treats undefined values as missing information; there are no RDF triples for those particular individuals or relationships. Details on the GlycoQL algorithm, semantic choices and the underlying system architecture are described in the Methods section.

## 3 Methods

### 3.1 Parsing algorithm

Let us recall that the GlycoCT encoding is defined as a connection table with a set terminology for monosaccharides and linkages. As such, this format allows for the unique identification of a glycan structure. The analysis of the GlycoCT syntax has already facilitated the GlySTreeM model design (Hayes *et al*. (2021)) and the data mapping algorithm that produces an RDF representation of the glycan tree structure. This tree, i.e., the structural pattern to search, requires additional processing to be translated into a SPARQL query.

The GlySTreeM pipeline parses structures expressed in GlycoCT into GlySTreeM RDF individuals. In this process, the data contained in the GlycoCT strings is used to build GlySTreeM knowledge. The GlycoCT syntax provides features to express the lack of information of some properties of the structures, often referred to as *unknowns*. These unknowns can be the anomeric configuration, the identification of the carbon atom on either the parent or child molecule or even the identity of the monosaccharide in question. This is illustrated in Figure 1 where the bottom right structure lacks information on monosaccharide identity (colourless shapes) and the top left one misses details of many linkages.

The unknowns are treated as information decorating the structure in the data parsing pipeline, but should not at all be considered as wild cards to preserve the semantic meaning of the pattern to search. For this reason, the unknowns in GlycoQL are considered as lack of knowledge hence the corresponding RDF triples are not included in the pattern derived from the GlycoCT string.

The pattern obtained allows retrieval of matching graphs, from structures with unknowns up to structures that are fully defined. This procedure is designed to respect the meaning of lack of knowledge that is key in our application field. In that way, the evaluation of the triples to include represents a semantic and substantial difference with respect to the GlySTreeM data mapping process as well as the entry point of the GlycoQL pipeline.

The next step involves the parsing of a search pattern in order to shape it into a suitable query that should result in a path through the GlySTreeM model, Figure 2. Whether this path provides an efficient SPARQL query is the remaining question. To answer it, the GlySTreeM model can be analysed to simulate possible outcomes. The representation of the glycan structure based on rigid parent-child relationships would imply traversing each structure from the root. This practice would be reasonable when the substructure to search is directly linked to the root of the structure tree (or containing it), such as a given core search (glycan cores are categorised and these categories are often searched; see Varki *et al*. (2015)). However, when searching for a substructure that might be in the middle or at the end of the tree (which makes sense biologically as many glycan substructures are known ligands), this approach would result in more computationally expensive and potentially inefficient queries. To avoid this situation, the representation of the glycan structure needs to be adjusted. To this end, the GlySTreeM model was extended with an additional representation of the monosaccharides (also called residues) composing the glycan structure (the nodes of the tree). This extension provides a flat (unstructured) representation of the residues so that each of them can serve as an entry point to navigate a specific tree, allowing the query engine to target residues at any level in the structure with equal complexity cost and then take advantage of the child-parent relationships to complete the search. The proposed representation requires grouping all the structure residues into a bag, called *ResidueSet* as shown in Figure 2, and this bag is linked to the Glycan instance.

**Fig. 2:**
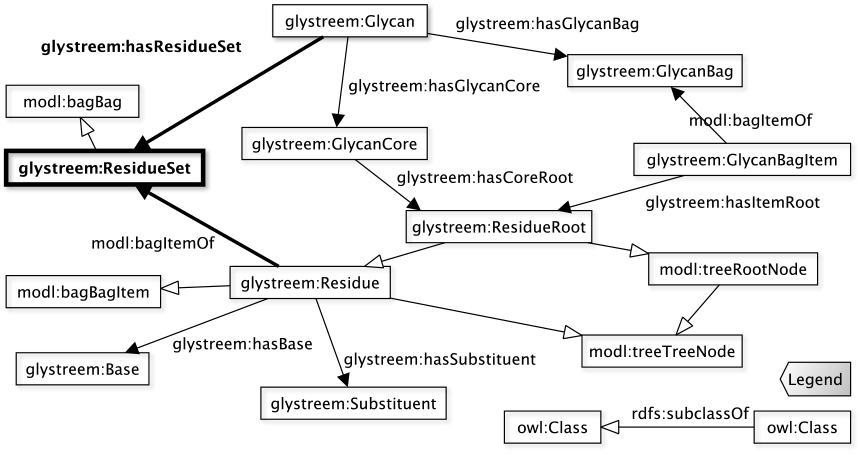
The ResidueSet class and the relationships (in bold) extending the GlySTreeM model.

The queries produced according to this model extension access the monosaccharides (residues) from the ResidueSets of each Glycan instance rather than from their tree structure root nodes. To increase the flexibility of the search two additional features have been added to the query: the possibility to freeze the total number of residues of the structure and the possibility to pin the substructure to query at the beginning (top/core) of the glycan.

Both features are implemented using two flags, respectively *ResNum* to set the number of the residues in the results to the one of the queried substructure; and *hasRoot* to specify that the first residue of the queried substructure is the *ResidueRoot*, hence at the first for all the retrieved structures.

### 3.2 Implementation

The GlySTreeM design guided the architecture that builds the SPARQL queries in GlycoQL, in particular the implementation relied on a similar technology stack enhanced with the integration to cover the SPARQL conversion. The parsing algorithm incorporating the semantic choices proper to GlycoQL is implemented as a Python module based on RdfLib (Krech (2006)). The module takes as input a GlycoCT string with the two flags *ResNum* and *hasRoot*, it produces a GlySTreeM pattern and translates it into a SPARQL query. To produce SPARQL syntax programmatically the algorithm uses the *SparqlBurger* library (Mitzias (2021)), a Python library that allows reproduction of several SPARQL constructs. The library has been then extended to produce other constructs not originally available such as “FILTER NOT EXISTS” and “HAVING”; the HAVING construct in particular has been instrumental to implement the logic for the *ResNum* flag.

The module implementing the parsing algorithm is embedded in the Flask Python framework (https://palletsprojects.com/p/flask/) to incapsulate it into a web based REST API service. GlycoQL users access this service using an HTML/javascript graphical user interface from their web browser. The web page allows the input of the requested parameters: GlycoCT in a dedicated textfield and *ResNum* and *hasRoot* as boolean checkboxes. After the submission the parameters are sent to the GlycoQL service to produce the SPARQL query, which is then forwarded to the triples store. The response is finally sent to the user interface and processed in the web page for rendering, including the depiction of the submitted pattern and the resulting structures shown in the SNFG format. The service architecture is described in Figure 3, while the GlycoQL service is available at https://glyconnect.expasy.org/glycoql/.

**Fig. 3:**
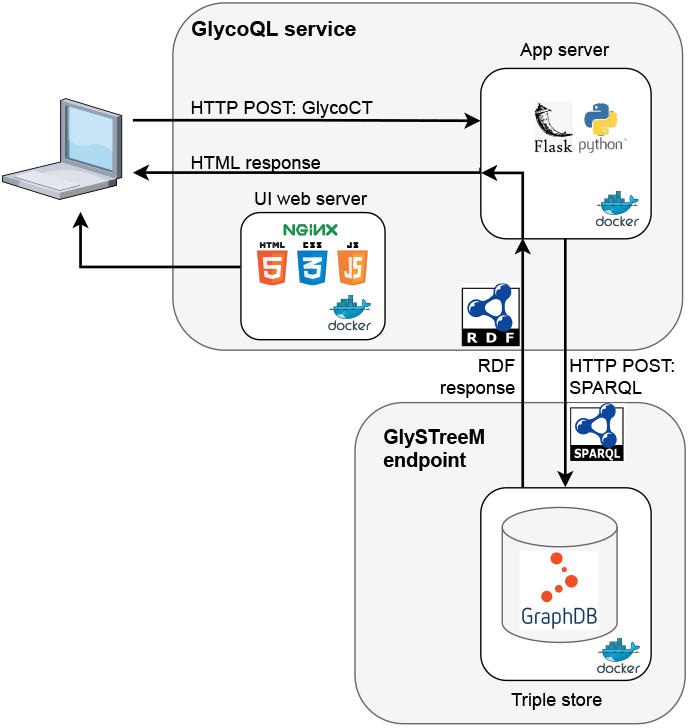
GlycoQL service architecture

### 3.3 Use Case - COVID dataset

The importance of glycosylation in the SARS-CoV-2 host-pathogen interaction has been shown in a number of recent papers (Gstöttner *et al*. (2021), Zhang *et al*. (2021), Zhou *et al*. (2021)), including N-linked glycosylation sites found on the viral surface spike protein, involved in host-cell entry. These references were included in a dedicated COVID section of GlyConnect. In this section, users can browse glycan structures that have been collated for the spike protein. These structures are reported for a number of glycosylation sites across experimentally different recombinant versions of the protein, using three expression systems, namely, HEK293 (human embryonic kidney), BTI-Tn-5B1-4 (insect cell) and CHO (Chinese Hamster Ovary). It appears useful to represent the data in the form of structural clusters that more accurately portray the profile of each site and/or recombinant protein.

Federated SPARQL queries were used to identify N-linked glycosylation sites and their associated glycan structures from UniProt, (Consortium (2021)), GlyConnect and GlySTreeM. These structures were manually inspected and consensus patterns were encoded using GlycoCT. GlycoQL was used to translate these encodings into queries, which were implemented in a semi-automated pipeline to classify glycan structures, Table 1. The queries are published on the GlySTreeM wiki page: https://glyconnect.expasy.org/glystreem/wiki.

**Table 1.** Use Case-COVID Dataset

|  |  |
| --- | --- |
| **Description** | Investigate differences between glycosylation sites, whole proteins and disease state. |
| **Actor** | Bioinformatician/Scientist. |
| **Initial Conditions** | GlyConnect database/GlySTreeM triple store<sup>1</sup>. |
| **Actor Actions** | **Systems Response** |
| 1) Federated SPARQL query for N-linked glycosylation sites(UniProt) and glycan IDs(GlyConnect) for COVID spike protein. | 2) 14 sites with 1035 associated glycans (both structures and compositions) on 4 recombinant spike proteins (UniProt P0DTC2). |
| 3) GlySTreeM query for glycans with associated GlycoCT | 4) 370 associated glycans (128 unique glycan structures.<sup>2</sup> |
| 5) Manual inspection of glycans. | 6) 5 GlycoCT encoded consensus patterns. |
| 7) GlycoQL translation of GlycoCT patterns | 8) Pipeline to automate glycan clustering. |
| 8) Query glycan list with GlycoQL patterns. | 9) Output of pattern distribution for entire set, for each glycosylation site, and for each site on each of four proteins. |
| **Post Conditions** |  |
| The results show the difference between glycosylation distribution when displayed with individual glycans versus glycan patterns. As the pattern matching was semi-automated, it was possible to easily create the distributions across sites, and individual recombinant proteins. |  |

<sup>1</sup> Snapshot is shown in Figure 4a; <sup>2</sup> See Figure 4b for distribution

## 4 Discussion

### 4.1 Development of GlycoQL

The design of GlySTreeM has opened many perspectives through which glycan structures can be analysed with the goal of comparing glycomic profiles whether associated with a protein glycosite, a glycoprotein, a cell line or a tissue in one or more condition(s). The resulting new prospects are being explored by shaping diversely complex queries. We suggested to rely on UniProt, GlyConnect and GlySTreeM to begin with, but such queries can also include other sources (federated) to broaden the scope of use cases and increase the coverage of integrated information. In order to approach these use cases and thus assess the potential of the GlySTreeM model, it became necessary to define a more direct way of access through which the main entities of these use cases could be refined with all possible features at no extra cost in complexity. GlycoQL was conceived out of the need to fill a gap in complexity and accessibility that the SPARQL language alone could not bridge. The use cases discussed first in the Methods section, and then below, bear witness to the fact that this project is aimed at facilitating everyday research work in glycobiology.

### 4.2 Use Case - COVID Dataset

A semi-automated classification pipeline was developed, using GlycoQL translations of GlycoCT encoded glycan consensus patterns, Table 1. The test set (identified using federated queries across three SPARQL endpoints), consisted of 370 (128 unique) glycan structures across 14 sites, described on four recombinant SARS-CoV-2 spike proteins, Figure 4a.

**Fig. 4:**
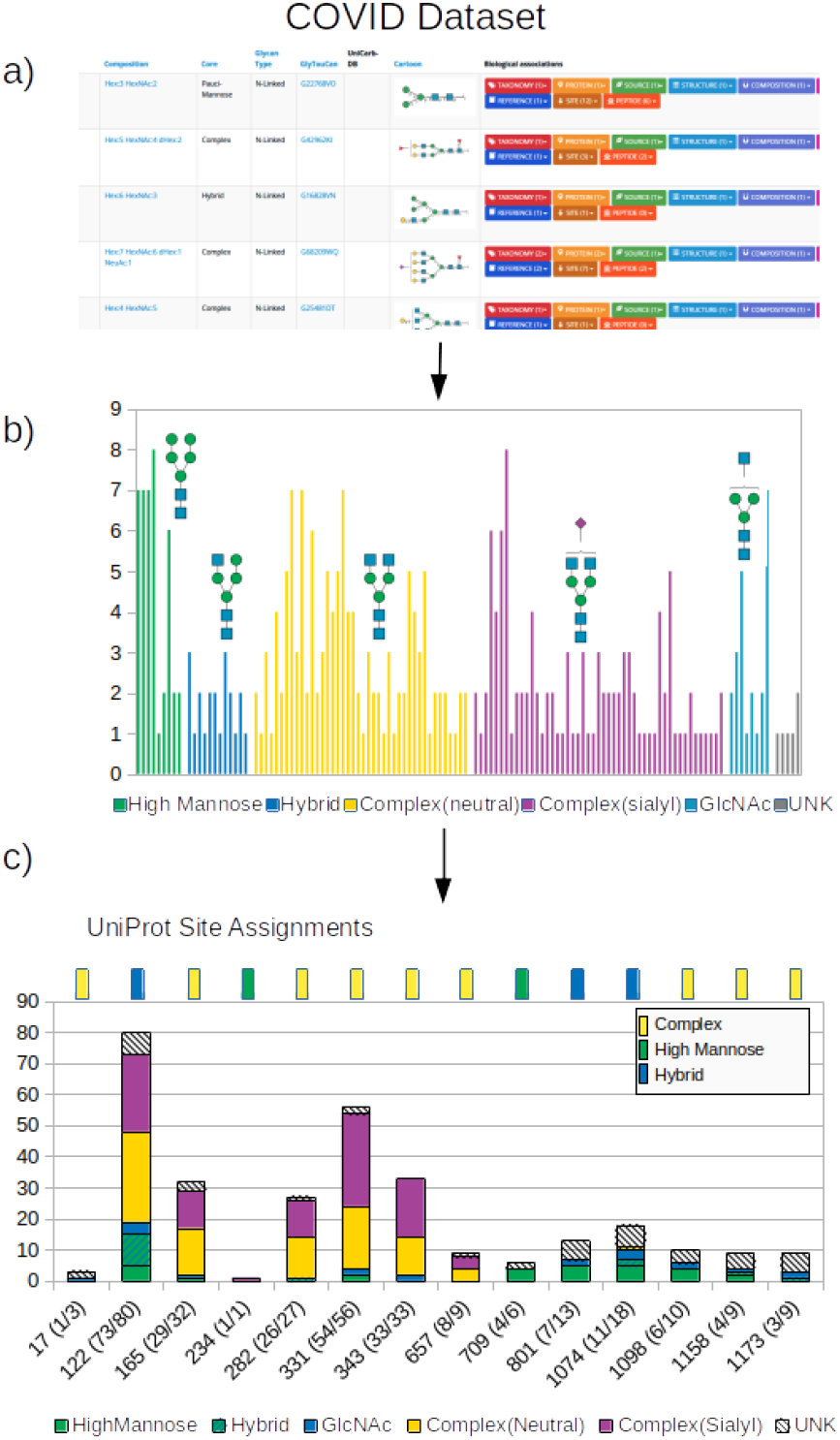
a) COVID dataset (GlyConnect), showing a variety of the glycan structures on various sites for four different recombinant spike proteins. b) a count profile of each of the glycans across all 14 sites on four recombinant proteins. GlycoQL patterns are shown as SNFG cartoons. c) The set of glycan structures was clustered using GlycoQL patterns, and this allowed a breakdown of the distribution within each glycosylation site. On top of the bar chart is the glycan type assignment as given in UniProt. Site number is given on the x axis. Numbers in brackets refer to the total number of glycans assigned a pattern/total number of glycans per site

Initially these 128 unique glycans were graphed to show counts for each structure across 14 sites and four recombinant spike proteins, Figure 4b. The glycans were manually annotated into types, which describe the main N-linked glycans (Varki *et al*. (2015)); high mannose, hybrid, neutral complex, sialylated complex and mono-GlcNAc extension. Fourteen glycans did not fit a pattern, and represent truncated or undefined structures. The five consensus patterns were generated as GlycoCT encodings, shown as SNFG cartoons, Figure 4b, which were translated into SPARQL queries with GlycoQL. The semi-automated classification pipeline was implemented to cluster the glycans according to these patterns which allowed a profile to be built for each glycosylation site, Figure 4c. The graph highlights the most heterogenous sites with respect to glycan structure, and also gives an indication as to the most abundant N-linked glycan type found at that location. This is compared to the UniProt assignment to the site (found in the PTM section of the protein entry, P0DTC2), shown as rectangles at the top of Figure 4c. Overall, the major trends reported in UniProt are confirmed, yet more detailed in our results. In particular, the important impact of sialylation can readily be spotted.

Furthermore, the most densely glycosylated site at position 122 displays a variety of properties impossible to report in UniProt annotations that assign site 122 as containing hybrid type structures. Our structural classification indicates that the profile is predominantly complex.

While an overall profile of the sites is an instant snapshot of site heterogeneity, the data represents four recombinant proteins, either full or partial sequence, and from three different expression systems. A site profile for each of the four was generated using the GlycoQL pipeline. Figure 5a and b represent full-length proteins, whereas Figure 5c and d, are the receptor-binding domain of the protein, amino acid 437 to 508. The glycosylation profile of the insect cell, Figure 5b, is obviously very different to that of either HEK (Figure 5a and c) or CHO (Figure 5d) cells. It is also evident that the majority of the undefined glycans are members of this profile. There is only one site that is effectively glycosylated with complex type structures, site 234. However the site profiles of 331 and 343 (RBD domain) are high mannose or undefined. This is in contrast to the other three profiles. All show an abundance of complex type structures and few truncated type glycans in the RBD domain. Given the role of the RDB domain in the virus-host interaction and considering that receptor binding is potentially hindered by glycans, then their structural details are likely to shed light on this interference. Furthermore, these results also can help assess the impact of the expression system.

**Fig. 5:**
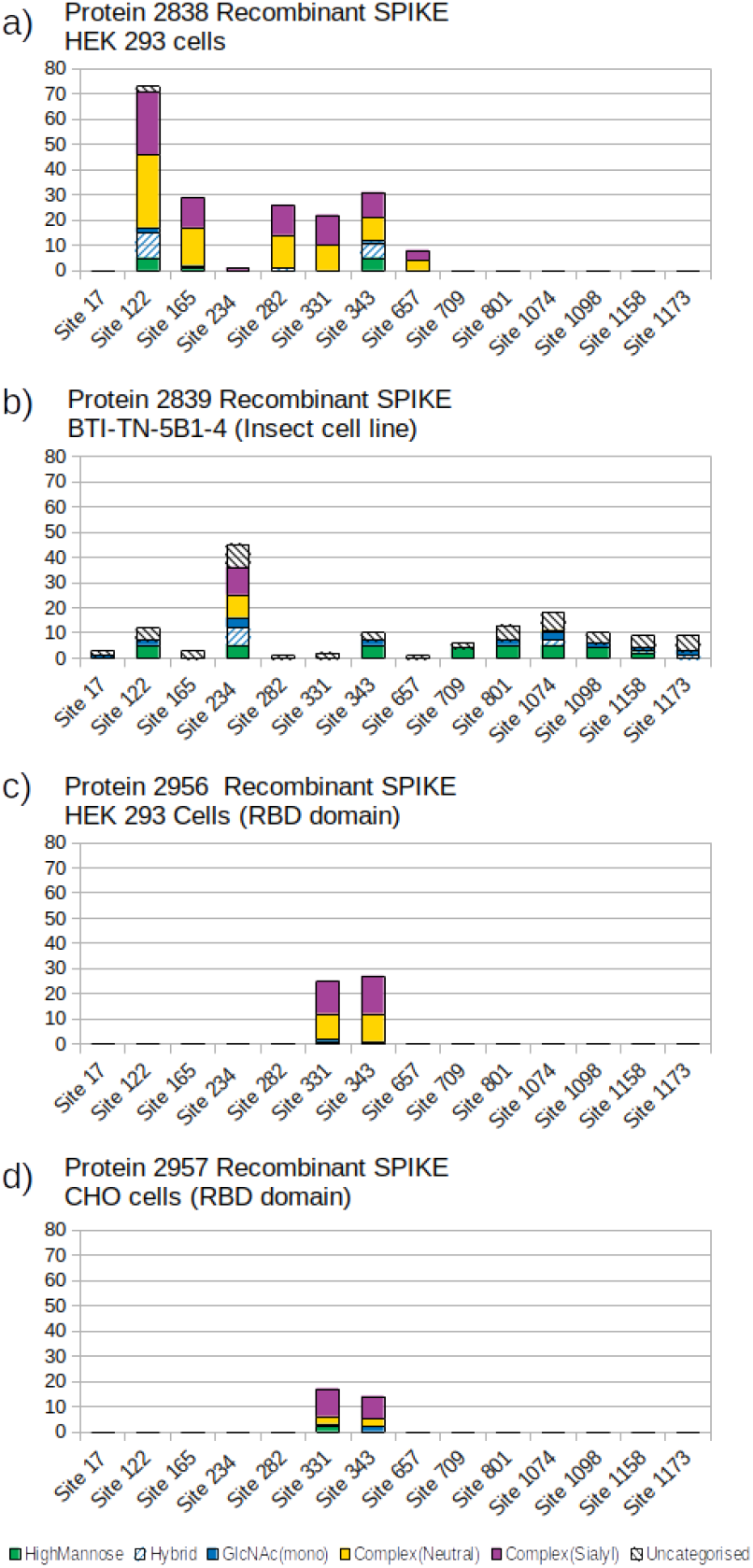
Site specific glycosylation profiles for four recombinant COVID spike proteins. The protein number refers to the GlyConnect Protein ID. Profile a) and b) represent the full AA sequence of the spike protein whereas profiles c) and d) consist of the receptor binding domain (RBD), amino acid 437 to 508. Sequence details can be found at https://www.uniprot.org/uniprot/P0DTC2.

The above use-case reflects the potential for this approach to evaluate and display the glycomic profile of proteins. The ability to reduce glycan structure complexity while preserving site heterogeneity will enhance available datasets in GlyConnect. It also gives a more realistic picture of site profiles as evidenced by the difference in site type assignments between our approach and UniProt.

## 5 Conclusion

The creation of the GlySTreeM triple store facilitated navigation of the structural space of individual glycans. This added value led to further development of tools to allow profiling of glycan sets. Here we present this approach as applied to the currently relevant spike protein from SARS-CoV-2. This profiling exercise allows us to quickly classify glycans using GlycoQL-derived patterns and display these according to site. These patterns describe N-linked glycosylation and can be used for other data sets, based on proteins, tissue, disease state etc.

The integration of such a pipeline into the GlyConnect/GlySTreeM family will create added-value knowledge, that better represents the underlying data and allow scientific insights that were not possible by scanning lists of individual glycan structures as often presented in glycomics or glycoproteomics related publications and associated datasets.

